# CPSM: R-package of an Automated Machine Learning Pipeline for Predicting the Survival Probability of Single Cancer Patient

**DOI:** 10.1101/2024.11.14.623597

**Authors:** Harpreet Kaur, Pijush Das, Kevin Camphausen, Uma Shankavaram

## Abstract

Accurate survival prediction is vital for optimizing treatment strategies in clinical practice. The advent of high-throughput multi-omics data and computational methods has enabled machine learning (ML) models for survival analysis. However, handling high-dimensional omics data remains challenging.

This study introduces the Cancer Patient Survival Model (CPSM), an R package developed to provide individualized survival predictions through a fully integrated and reproducible computational pipeline. The CPSM package encompasses nine modules that streamline the survival modeling workflow, organized into four key stages: (1) Data Preprocessing and Normalization, (2) Feature Selection, (3) Survival Prediction Model Development, and (4) Visualization. The visual tools facilitate the interpretation of survival predictions, enhancing clinical decision-making. By providing an end-to-end solution for multi-omics data integration and analysis, CPSM not only enhances the precision of survival predictions but also aids in discovering clinically relevant biomarkers.

**Availability and Implementation:** The CPSM Package is freely available at the GitHub URL: https://github.com/hks5august/CPSM

## 1. Introduction

In the current era, predicting cancer patient survival using multi-omics data has become a crucial focus of biomedical research, as it plays a key role in therapeutic interventions and patient management (Abbasi, et al., 2024; Bashiri, et al., 2017; Cheon, et al., 2016). The rapid development of RNA sequencing technology has led to an extensive generation of multi-omics profiles from cancer patients. With the increasing availability of these datasets and advancements in computational methods, a wide array of user-friendly primary and secondary data resources has been developed. These resources enable researchers to explore clinical and omics data to identify diagnostic and prognostic biomarkers across various cancer types (Barretina, et al., 2012; Cai, et al., 2019; Cancer Genome Atlas Research, et al., 2013; Cerami, et al., 2012; Deng, et al., 2017; Deng, et al., 2023; Grossman, et al., 2016; Joly, et al., 2012; Kaur, et al., 2020; Rhodes, et al., 2004; Tate, et al., 2019; Uhlen, et al., 2005; Yang, et al., 2019; Zhao, et al., 2021).

The surge in multi-omics data availability has profoundly impacted the field of cancer research, providing unprecedented opportunities for biomarker discovery and precision medicine (Bhalla, et al., 2017; Bhalla, et al., 2019; Deng, et al., 2023; Dhall, et al., 2020; Garg, et al., 2024; Han, et al., 2013; Hussein, Abou-Shanab and Badr, 2024; Kaur, Bhalla and Raghava, 2019; Kaur, et al., 2019; Oh, et al., 2020; Vasudevan and Murugesan, 2018; Xiao, et al., 2021; Xiao, et al., 2022; Yang, et al., 2024; Zhang and Liu, 2021). However, the inherent complexity and heterogeneity of multi-omics data pose significant challenges for analysis, interpretation, and subsequent biomarker identification (Brooks, et al., 2024; Lopez de Maturana, et al., 2019; Matthews, Hanison and Nirmalan, 2016; McDermott, et al., 2013; Mohr, et al., 2024; Ng, et al., 2023; Yamada, et al., 2021). Extracting meaningful biological insights from these data requires a series of rigorous computational steps, each demanding specific expertise in areas such as data pre-processing, normalization, statistical analysis, feature selection, predictive modeling, and visualization.

Although numerous specialized tools exist to address individual analytical tasks, that are often designed to function independently (Biecek, 2021; Bolstad, 2024; Gerds, 2023; Haider, 2019; Kuhn, 2008; Sing, et al., 2005; Wickham, 2016). The lack of integration necessitates the use of multiple workflows, which can present considerable challenges, especially for researchers lacking extensive computational expertise. Additionally, the inherent heterogeneity of multi-omics data further complicates the seamless application of these tools, frequently resulting in inefficiencies and creating significant barriers to effective analysis.

To address these challenges, we proposed a comprehensive computational pipeline that integrates and customizes a range of computational tools into a cohesive and user-friendly workflow. This pipeline is specifically designed to facilitate survival prediction in cancer patients by leveraging both clinical and gene expression data. By streamlining the multi-step analytical process into a unified framework, our approach seeks to enhance the precision of survival predictions and support the identification of clinically relevant biomarkers, thereby contributing to the advancement of personalized oncology. We illustrate its use by showcasing its functionality in two use cases employing publicly available data from The Cancer Genome Atlas Program (TCGA) for two malignancies, Glioblastoma (GBM) and Kidney Renal Clear Cell Carcinoma (KIRC).

## 2. Algorithms and Modules

The CPSM package comprises nine key modules/functions, each designed for a specific task in the survival analysis workflow. These tasks span the entire modeling pipeline, from data preprocessing, splitting data into training and test sets, data normalization, feature selection, survival prediction model development, validation, visualization, and nomogram generation, as outlined in Figure 1. These modules can be categorized into four main stages: (1) Data Preprocessing and Formatting, (2) Feature Selection, (3) Prediction Model Development and (4) Visualization. Each step, along with the functions from the CPSM package that corresponds to it, are detailed below.

**Figure 1:**
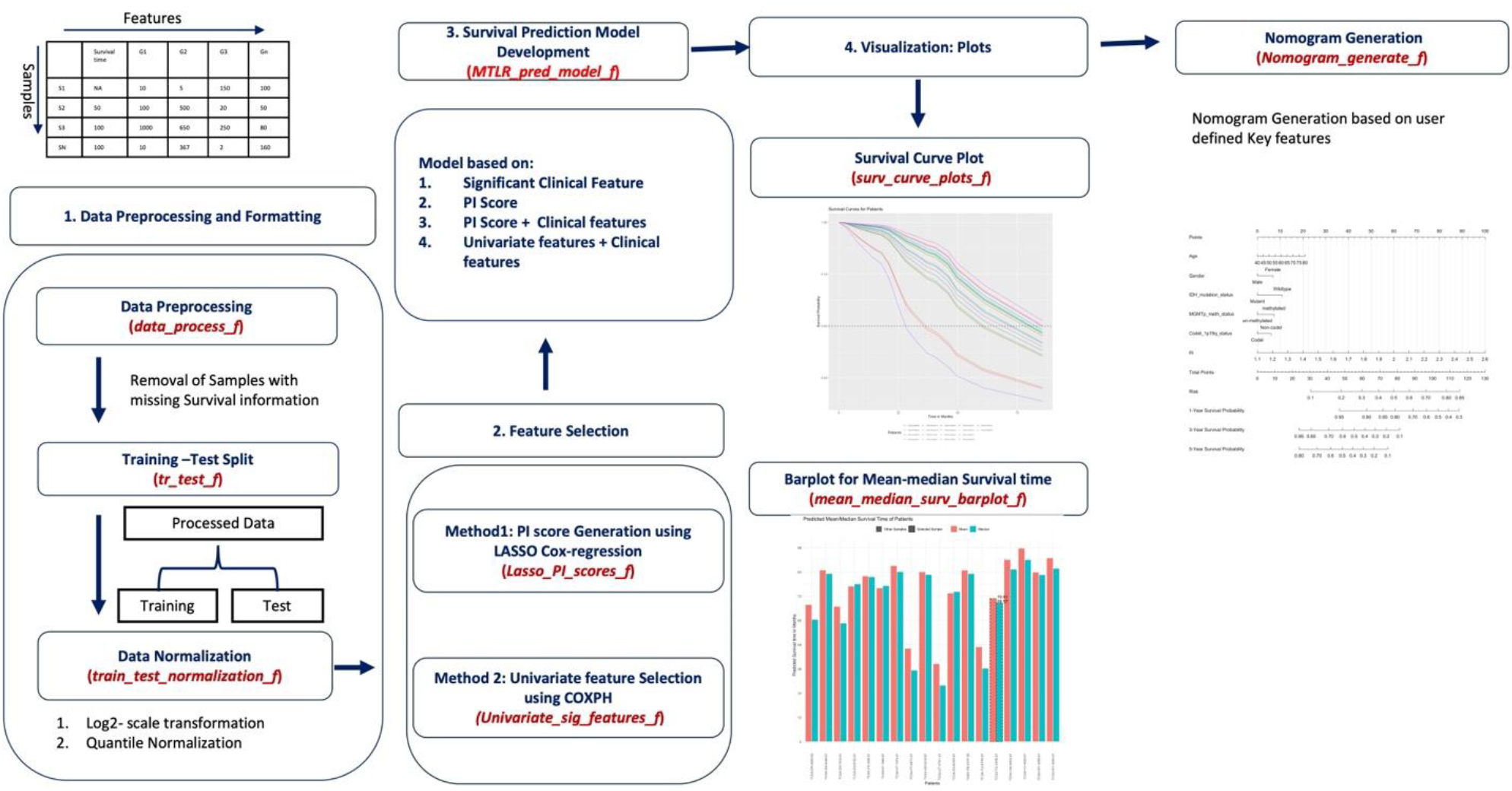
Overall structure and workflow of the CPSM (Cancer Patient Survival Model) illustrating the 9 key modules employed in each step.

### 2.1 Data Preprocessing and Normalization

Effective data preprocessing is critical for reliable downstream analysis in survival prediction. Proper formatting, normalization, and splitting of data into training and test sets are prerequisites for model development. In CPSM, this step is critical specifically for high-throughput gene expression data and survival information. The 3 primary preprocessing functions in CPSM are described below.

#### 2.1.1 data_process_f

This function is responsible for converting survival times from days to months, if the input is presented in days. It also filters out samples with missing overall survival (OS) information. The expected input data typically contain gene expression data in formats such as FPKM, RPKM, TPM, or RSEM. The function appends a new column to the dataset, labeled “OS_month,” which serves as the survival-time feature in the data.

#### 2.1.2 tr_test_f

This function partitions the dataset into training and test sets for the purpose of model development and validation. Users can define the ratio of data split by providing a fraction value (e.g. 0.9 or 0.8 or 0.7) that will used to split data into training and testing sets. For instance, fraction=0.9 will split data into 90% training and 10% as the test set. This split ensures that feature selection and model development are performed on the training set while the test set remains untouched for unbiased model evaluation.

#### 2.1.3. train_test_normalization_f

Gene expression data often exhibit a high degree of variability across samples, necessitating normalization to ensure that features are comparable. This function applies two key transformations:

I. **Log-scale transformation**: Converts raw FPKM/RPKM/TPM/RSEM values to a logarithmic scale using the formula:

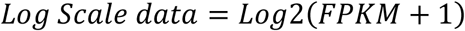 This mitigates the skewness of data distributions.
II. **Quantile Normalization** Quantile normalization is applied to both the training and test datasets, with the training set used as a reference matrix. This ensures consistency in the data distributions across samples, a critical step for machine learning algorithms.

These steps create a standardized, normalized dataset, prepared for feature selection and model training.

### 2.2 Feature Selection

The next step in the CPSM pipeline involves identifying features (typically genes) that are most relevant for predicting patient survival. Since high-dimensional gene expression data often contains noise or redundant features, this step helps reduce dimensionality and enhances the model’s performance. CPSM offers two distinct methods for feature selection, enabling users to select the option that best aligns with their specific requirements:

#### 2.1.1 Lasso_PI_scores_f

The Least Absolute Shrinkage and Selection Operator (LASSO) regression is a powerful technique for feature selection, especially in high-dimensional data. This function computes **Prognostic Index (PI) Scores** for each sample, which are a weighted sum of selected gene expression values, where the weights (beta coefficients) are determined via LASSO. Features with non-zero coefficients are retained. The PI score is computed based on the expression of the selected features and their beta coefficients as follows:

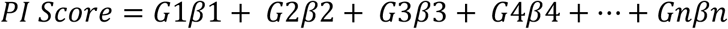

Where, G1, G2, G3, G4, and G5 are the genes or features selected by the LASSO method (G1 gene1, G2 - gene2, G3- gene3, G4-gene4, Gn-nth gene) and β1, β2, β3, β4, βn are their corresponding beta coefficient values, respectively. *Lasso_PI_scores_f* function will provide the following outputs: (1)A list of selected features with corresponding beta coefficients. (2) A lambda plot from LASSO regression, showing the penalty applied to coefficients as the regularization parameter increases. (3) Training and test datasets appended with the PI score for each sample.

#### 2.1.2 Univariate_sig_features_f

This function performs univariate **Cox Proportional Hazards (CoxPH)** regression for each feature in the dataset. The goal is to identify features that are individually associated with survival outcomes, based on a significance threshold (commonly, p < 0.05). The output includes: (1) Coefficients for each significant feature. (2) Hazard Ratios (HR) indicating the impact of each feature on survival. (3) corresponding p-values and concordance index. (4) Training and test datasets containing significant features, and associated survival metrics.

Together, these two methods enable users to select relevant features and focus on those that contribute most to the prognostic model, thus improving its interpretability and predictive accuracy.

### 2.3 Survival Prediction Model Development

Once the relevant features are identified, CPSM provides tools to develop robust prediction models and visualize their results. The model primarily employed is based on Multi-Task Logistic Regression (MTLR), which is well-suited for survival analysis in high-dimensional data settings. This MTLR-based approach allows for dynamic survival predictions that are not restricted to fixed time intervals, unlike traditional CoxPH models, providing more flexibility and improved accuracy. The main functions for model development is described below:

#### MTLR_pred_model_f

Prediction function is built using the selected features and can be customized to develop models based on different types of features available, if the user has only clinical features, both gene expression and clinical data, or only gene expression data, respective models are made by — clinical features alone, the PI score alone, PI score with clinical features and clinical features with gene expression of univariate features. Each Model with its utility is explained in detail in supplementary material section 1. The key outputs of this function are: (1) A training model, (2) A table with predicted survival probabilities for each patient at multiple time points, (3) A table containing predicted mean and median survival times for individual patients, (4) A table presenting the performance metrics, including the Integrated Brier Score (IBS) and the Concordance Index (C-Index), for both training and test datasets based on the prediction model. These metrics range from 0 to 1, with lower IBS values and higher C-Index values indicating better model performance, while higher IBS and lower C-Index values suggest poor model performance.

### 2.4. Visualization

The CPSM package includes a range of visualization tools to enhance the interpretation of survival predictions and support clinical decision-making. These functions enable users to explore survival probabilities, compare predicted survival times, and assess individual risk factors through intuitive visualizations like survival curves, bar plots, and nomograms. The key functions designed for visualization are described below:

#### 2.4.1 surv_curve_plots_f

For intuitive visualization, CPSM package allows user to check survival probability of each sample of the test data in the form survival curves. This function generates plots based on the survCurves_data that was obtained from the *MTLR_pred_model_f* function. Besides, this function allows user to visualize a specific patient on the curve.

#### 2.4.2 mean_median_surv_barplot_f

This function allows user to visualize predicted median and mean survival of samples of test data on the bar plots. This function generates bar plots based on the data containing mean and medium information, which was obtained from the *MTLR_pred_model_f* function Besides, this function also allows to visualize a specific patient on the plot.

#### 2.4.3 Nomogram_generate_f

Nomogram is two-dimensional graphical visualization tool used in clinical settings. It is designed by integrating patient characteristics with statistical or mathematical models used to estimate outcome of patients. It is commonly used to predict risks like survival, cancer recurrence, or metastasis, aiding in personalized treatment decisions and patient counseling (Kattan and Marasco, 2010; Wingyi Lee, 2022). The *Nomogram_generate_f* function of CPSM will allow users to generate a nomogram plot for their data (training data containing all samples) based on user-defined clinical and other features in their data. The generated nomogram can estimate individual risks, such as the probability of survival at 1, 3, 5, or 10 years, and is particularly useful for clinical decision-making.

## 3 Implementation

The *CPSM* has been developed with R version 4.3.3 (Team, 2024). It integrates various R-libraries including “caret” (Kuhn, 2008), “preprocessCore” (Bolstad, 2024), “survival”, “survminer” “glmnet” (Friedman, Hastie and Tibshirani, 2010; Simon, et al., 2011), “ggplot2” (Wickham, 2016), “SurvMetrics” (Wang, 2022), “MTLR” (Haider, 2019), “rms” (FE, 2024), “survival” (T, 2024), “survminer” (Biecek, 2021), “survivalROC” (Saha-Chaudhuri, 2022), “pec”, etc. A description of the instructions on installing and running the *CPSM* is present on the public Github repository along with an example dataset. A detailed workflow for each step and parameters is also provided in the detailed manual on the GitHub page.

## 4 Case studies

To explore the utility of the CPSM package, we conducted analyses on two case studies obtained from The Cancer Genome Atlas (TCGA) for Glioblastoma (GBM) and Kidney Renal Clear Cell Carcinoma (KIRC). These datasets comprise RNAseq gene expression data for 38,693 transcripts represented as FPKM values, along with 19 clinical and demographic features (Table S1), including overall survival (OS) information. Our primary objectives were to identify significant prognostic features and develop robust survival models for predicting patient survival probabilities using the CPSM package. Detailed methodologies and results for each case study can be found in the supplementary materials. For each case study, we tested all four training models using clinical features alone (Model 1), the PI score alone (Model 2), PI score with clinical features (Model 3), and clinical features plus gene expression of univariate features (Model 4). Below, we summarize the key findings from each analysis.

### 4.1 Case Study1

In the case of TCGA-GBM, Model 2, derived from the Prognostic Index (PI) score, demonstrated superior performance on training data of 144 samples with a C-Index of 0.80 and an IBS of 0.068 (Supplementary Table S2). The PI score was generated using 36 RNA features selected through the **Lasso_PI_scores_f** function of the CPSM package. On a test cohort of 17 samples, it achieved a C-Index of 0.66 and an integrated IBS of 0.083. For the demonstration, we have chosen a specific patient with barcode ID “TCGA-41-2572-01” with an overall survival (OS) time of 13 months. Model 2 predicted a median survival time of 13.3 months (within 5% error range) for this patient (Supplementary Figure S2A). Since the PI score-based model demonstrated the best performance, we developed a nomogram (Figure S3) using the **Nomogram_generate_f** function from the CPSM package. This nomogram, derived from the PI scores of the training data, was designed to predict patients’ 1-year, 3-year, and 5-year survival risks, achieving a C-Index of 0.72.

### 4.2 Case Study2

In the case of TCGA-KIRC, Model 3, which integrated three clinical features (age, subtype, and pathological grade) with the PI score, yielded the highest predictive performance on training data of 473 samples with a C-Index of 0.79 and IBS score of 0.16. Notably, the PI score in this model was constructed from 9 RNA features (Table S5) identified using the **Lasso_PI_scores_f** function. Model 3 remains superior performer on test dataset of 53 samples and attained a C-Index of 0.76 and an IBS of 0.23 (Supplementary Table S4). For testing we randomly selected a patient with TCGA barcode ID “TCGA-CJ-6030-01” with overall survival (OS) time of 76 months. The model predicted a median survival time of 73.81 months (within 5% error range) for this patient (Supplementary Figure S5A). Given the strong predictive capability of Model 3 in the TCGA-KIRC data, we developed a nomogram (Figure S6) based on the PI score, age, subtype, and pathological grade to estimate 1-year, 3-year, 5-year, and 10-year survival probabilities, which achieved a C-Index of 0.76.

## Discussion and Conclusion

The identification of pertinent prognostic signatures for survival prediction is crucial for optimizing treatment strategies in cancer management. CPSM is specifically designed to predict cancer patient survival by integrating clinical and gene expression data. By consolidating the multi-step analytical process into a unified framework, CPSM simplifies the analytical workflow and facilitates the discovery of clinically relevant biomarkers. This approach aims to enhance the precision and interpretability of survival predictions, ultimately supporting both research efforts and potential clinical applications in personalized medicine.

The CPSM represents a comprehensive and automated survival prediction pipeline, aimed at expanding accessibility to advanced survival analysis by reducing the dependency on specialized programming skills. By integrating a range of established survival associated tools in a customized manner, CPSM enables users to effectively utilize survival-relevant features within a customizable and multifunctional framework and generating Prognostic Index (PI) scores for individual samples, respectively. The customized incorporation of the MTLR (Haider, 2019) significantly augments its utility, facilitating the development of robust survival probability prediction models utilizing selected survival relevant feature sets from complex, high-dimensional datasets. Importantly, the survival-associated features identified through CPSM’s feature selection module not only contribute to predictive accuracy of the model but also hold potential as independent biomarkers, warranting further exploration in clinical research.

Although CPSM automates many steps in the analytical workflow, it maintains considerable flexibility through its range of customized options. This allows users to design models according to specific feature sets derived from the package’s feature selection modules, whether these are significant clinical features, PI score, or combination of them and computation of clinical and molecular features identified through univariate survival analysis. Importantly, CPSM’s applicability is not restricted to clinical or transcriptomic data but extends to a wide variety of quantitative data types, including proteomic and metabolomic datasets. This versatility broadens the relevance of CPSM across diverse research domains beyond transcriptomics oncology.

CPSM employs a suite of robust R packages that ease the processes of data preprocessing, normalization, feature selection, and model development. The tools “caret” and “preprocessCore” (Bolstad, 2024; Kuhn, 2008) support efficient data preprocessing and normalization, while packages like survival, survminer, and glmnet (Biecek, 2021; Friedman, Hastie and Tibshirani, 2010; T, 2024) enable effective feature selection and prioritization. Additionally, CPSM leverages SurvMetrics, MTLR, and rms (FE, 2024; Haider, 2019; Saha-Chaudhuri, 2022; Wang, 2022) for survival prediction model development and the creation of nomograms, facilitating a seamless user experience that minimizes the need for extensive coding expertise. This approach also addresses common formatting issues associated with input and output, further enhancing accessibility for non-bioinformaticians.

When compared to other survival prediction tools, such as COXPH, glmnet, and Neural Multi-Task Logistic Regression (N-MTLR) (Fotso, 2018; Simon, et al., 2011; T, 2024; Terry M. Therneau, 2000), CPSM offers unique functionalities specifically designed for survival analysis. CPSM calculates p-values, hazard ratios (HR), beta coefficients, and the concordance index (C-Index) for individual features, with automatic selection of statistically significant features (p < 0.05). Moreover, CPSM computes PI scores for each sample in both training and testing datasets, which enhances the reliability of survival predictions. Unlike traditional Cox models, CPSM employs MTLR to estimate survival probabilities over multiple time intervals, providing a comprehensive framework for clinical decision-making. Furthermore, CPSM extends the capabilities of MTLR by offering survival probability predictions alongside mean and median survival times, accompanied by graphical representations such as survival curves and bar plots, which collectively enrich the interpretability of results.

The accessibility of CPSM is a significant advantage, as it does not require extensive coding expertise. The diverse functionalities of CPSM apart it from other survival analysis packages, which often target specifically bioinformaticians with advanced R programming skills. The utility of CPSM has been demonstrated with real-world datasets. For instance, demonstrating using our case studies, in the TCGA-GBM cohort, a model based on PI scores (Model-2) demonstrated superior performance relative to other models based on other selected feature sets, while in the TCGA-KIRC cohort, a model combining clinical features and PI scores (Model-3) showed improved predictive accuracy. Conversely, models based on large numbers of univariate features, such as Model-4 in the TCGA-KIRC cohort, underperformed. These results underscore the importance of prioritizing features based on metrics like HR, C-Index, and p-values to achieve optimal model performance. Consequently, it is recommended that researchers carefully select univariate features based on these metrics to enhance the robustness and effectiveness of survival prediction models. One of the other key strengths of CPSM is its capability to generate nomograms, a feature often absents in other survival prediction tools. Nomograms provide a visual representation of the relationship between multiple prognostic factors and survival outcomes, offering a highly interpretable tool that supports clinicians in making informed, personalized treatment decisions.

In conclusion, the CPSM provides an adaptable, user-friendly, and scientifically rigorous solution for survival analysis and biomarker discovery in cancer research. Its ability to accommodate diverse data types, coupled with its advanced functionalities, such as the creation of nomograms, positions CPSM as an invaluable tool for researchers and clinicians in the field of cancer survival prediction. By integrating well-established methodologies into an accessible platform, CPSM holds significant potential to advance personalized cancer treatment and improve patient outcomes. Its utility extends beyond oncology, offering a versatile framework suitable for survival analysis across various biomedical research disciplines.

## Authors Contribution

HK and US designed the project. HK collected and created the datasets and developed the algorithms. HK and PD implemented the package. HK, PD, and US analyzed the results. US and KC coordinated the project. HK drafted and manuscript and PD and US refined the drafted manuscript. All of the authors have read and approved the final manuscript.

## Availability

The CPSM Package is freely available to download from the GitHub URL: https://github.com/hks5august/CPSM

## References

Abbasi, A.F., et al. Survival prediction landscape: an in-depth systematic literature review on activities, methods, tools, diseases, and databases. Front Artif Intell 2024;7:1428501.

Barretina, J., et al. The Cancer Cell Line Encyclopedia enables predictive modelling of anticancer drug sensitivity. Nature 2012;483(7391):603–603.

Bashiri, A., et al. Improving the Prediction of Survival in Cancer Patients by Using Machine Learning Techniques: Experience of Gene Expression Data: A Narrative Review. Iran J Public Health 2017;46(2):165–165.

Bhalla, S., et al. Gene expression-based biomarkers for discriminating early and late stage of clear cell renal cancer. Sci Rep 2017;7:44997.

Bhalla, S., Kaur, H., Dhall, A. and Raghava, G.P.S. Prediction and Analysis of Skin Cancer Progression using Genomics Profiles of Patients. Sci Rep 2019;9(1):15790.

Biecek, A.K.a.M.K.a.P. 2021. survminer: Drawing Survival Curves using ‘ggplot2. Release R package version 0.4.9. https://CRAN.R-project.org/package=survminer

Bolstad, B. 2024. preprocessCore: A collection of pre-processing functions. Release R package version 1.66.0. https://bioconductor.org/packages/preprocessCore

Brooks, T.G., Lahens, N.F., Mrcela, A. and Grant, G.R. Challenges and best practices in omics benchmarking. Nat Rev Genet 2024;25(5):326–326.

Cai, L., et al. LCE: an open web portal to explore gene expression and clinical associations in lung cancer. Oncogene 2019;38(14):2551–2551.

Cancer Genome Atlas Research, N., et al. The Cancer Genome Atlas Pan-Cancer analysis project. Nat Genet 2013;45(10):1113–1113.

Cerami, E., et al. The cBio cancer genomics portal: an open platform for exploring multidimensional cancer genomics data. Cancer Discov 2012;2(5):401–401.

Cheon, S., et al. The accuracy of clinicians’ predictions of survival in advanced cancer: a review. Ann Palliat Med 2016;5(1):22–22.

Deng, M., et al. FirebrowseR: an R client to the Broad Institute’s Firehose Pipeline. Database (Oxford) 2017;2017.

Deng, X., et al. Glioma-BioDP: database for visualization of molecular profiles to improve prognosis of brain cancer. BMC Med Genomics 2023;16(1):168.

Dhall, A., et al. Computing Skin Cutaneous Melanoma Outcome From the HLA-Alleles and Clinical Characteristics. Front Genet 2020;11:221.

FE, H.J. 2024. rms: Regression Modeling Strategies. Release R package version 6. 8–1. https://CRAN.R-project.org/package=rms

Fotso, S. Deep Neural Networks for Survival Analysis Based on a Multi-Task Framework. arXiv 2018.

Friedman, J., Hastie, T. and Tibshirani, R. Regularization Paths for Generalized Linear Models via Coordinate Descent. J Stat Softw 2010;33(1):1–1.

Garg, M., et al. Disease prediction with multi-omics and biomarkers empowers case-control genetic discoveries in the UK Biobank. Nat Genet 2024;56(9):1821–1821.

Grossman, R.L., et al. Toward a Shared Vision for Cancer Genomic Data. N Engl J Med 2016;375(12):1109–1109.

Haider, H. 2019. MTLR: Survival Prediction with Multi-Task Logistic Regression. Release R package version 0.2.1. https://CRAN.R-project.org/package=MTLR

Han, H., Li, X.L., Ng, S.K. and Ji, Z. Multi-resolution-test for consistent phenotype discrimination and biomarker discovery in translational bioinformatics. J Bioinform Comput Biol 2013;11(6):1343010.

Hussein, R., Abou-Shanab, A.M. and Badr, E. A multi-omics approach for biomarker discovery in neuroblastoma: a network-based framework. NPJ Syst Biol Appl 2024;10(1):52.

Joly, Y., et al. Data sharing in the post-genomic world: the experience of the International Cancer Genome Consortium (ICGC) Data Access Compliance Office (DACO). PLoS Comput Biol 2012;8(7):e1002549.

Kattan, M.W. and Marasco, J. What is a real nomogram? Semin Oncol 2010;37(1):23–23.

Kaur, H., Bhalla, S., Kaur, D. and Raghava, G.P. CancerLivER: a database of liver cancer gene expression resources and biomarkers. Database (Oxford) 2020;2020.

Kaur, H., Bhalla, S. and Raghava, G.P.S. Classification of early and late stage liver hepatocellular carcinoma patients from their genomics and epigenomics profiles. PLoS One 2019;14(9):e0221476.

Kaur, H., Dhall, A., Kumar, R. and Raghava, G.P.S. Identification of Platform-Independent Diagnostic Biomarker Panel for Hepatocellular Carcinoma Using Large-Scale Transcriptomics Data. Front Genet 2019;10:1306.

Kuhn, M. Building Predictive Models in R Using the caret Package. Journal of Statistical Software 2008;28(5):1–1.

Lopez de Maturana, E., et al. Challenges in the Integration of Omics and Non-Omics Data. Genes (Basel) 2019;10(3).

Matthews, H., Hanison, J. and Nirmalan, N. “Omics”-Informed Drug and Biomarker Discovery: Opportunities, Challenges and Future Perspectives. Proteomes 2016;4(3).

McDermott, J.E., et al. Challenges in Biomarker Discovery: Combining Expert Insights with Statistical Analysis of Complex Omics Data. Expert Opin Med Diagn 2013;7(1):37–37.

Mohr, A.E., et al. Navigating Challenges and Opportunities in Multi-Omics Integration for Personalized Healthcare. Biomedicines 2024;12(7).

Ng, S., Masarone, S., Watson, D. and Barnes, M.R. The benefits and pitfalls of machine learning for biomarker discovery. Cell Tissue Res 2023;394(1):17–17.

Oh, S., et al. From genome sequencing to the discovery of potential biomarkers in liver disease. BMB Rep 2020;53(6):299–299.

Rhodes, D.R., et al. ONCOMINE: a cancer microarray database and integrated data-mining platform. Neoplasia 2004;6(1):1–1.

Saha-Chaudhuri, P.J.H.a.P. 2022. survivalROC: Time-Dependent ROC Curve Estimation from Censored Survival Data. Release R package version 1.0.3.1. https://CRAN.R-project.org/package=survivalROC

Simon, N., Friedman, J., Hastie, T. and Tibshirani, R. Regularization Paths for Cox’s Proportional Hazards Model via Coordinate Descent. J Stat Softw 2011;39(5):1–1.

T, T. 2024. A Package for Survival Analysis in R. Release R package version 3. 7–0. https://CRAN.R-project.org/package=survival

Tate, J.G., et al. COSMIC: the Catalogue Of Somatic Mutations In Cancer. Nucleic Acids Res 2019;47(D1):D941–D947.

Team, R.C. 2024. R: A Language and Environment for Statistical Computing

Terry M. Therneau, P.M.G. Modeling Survival Data: Extending the Cox Model. Springer, New York; 2000.

Uhlen, M., et al. A human protein atlas for normal and cancer tissues based on antibody proteomics. Mol Cell Proteomics 2005;4(12):1920–1920.

Vasudevan, P. and Murugesan, T. Cancer Subtype Discovery Using Prognosis-Enhanced Neural Network Classifier in Multigenomic Data. Technol Cancer Res Treat 2018;17:1533033818790509.

Wang, H.Z.a.X.C.a.S.W.a.Y.Z.a.H. 2022. SurvMetrics: Predictive Evaluation Metrics in Survival Analysis. Release R package version 0.5.0. https://CRAN.R-project.org/package=SurvMetrics

Wickham, H. ggplot2: Elegant Graphics for Data Analysis. Springer-Verlag New York; 2016.

Wingyi Lee, S.-K.L., Yuanpeng Zhang, Ruijie Yang, Jing Cai. Review of methodological workflow, interpretation and limitations of nomogram application in cancer study. Radiation Medicine and Protection 2022;3(4):200–200.

Xiao, Q., et al. High-throughput proteomics and AI for cancer biomarker discovery. Adv Drug Deliv Rev 2021;176:113844.

Xiao, Y., Bi, M., Guo, H. and Li, M. Multi-omics approaches for biomarker discovery in early ovarian cancer diagnosis. EBioMedicine 2022;79:104001.

Yamada, R., et al. Interpretation of omics data analyses. J Hum Genet 2021;66(1):93–93.

Yang, Y., et al. GliomaDB: A Web Server for Integrating Glioma Omics Data and Interactive Analysis. Genomics Proteomics Bioinformatics 2019;17(4):465–465.

Yang, Z., Guan, F., Bronk, L. and Zhao, L. Multi-omics approaches for biomarker discovery in predicting the response of esophageal cancer to neoadjuvant therapy: A multidimensional perspective. Pharmacol Ther 2024;254:108591.

Zhang, Z. and Liu, Z.P. Robust biomarker discovery for hepatocellular carcinoma from high-throughput data by multiple feature selection methods. BMC Med Genomics 2021;14(Suppl 1):112.

Zhao, Z., et al. Chinese Glioma Genome Atlas (CGGA): A Comprehensive Resource with Functional Genomic Data from Chinese Glioma Patients. Genomics Proteomics Bioinformatics 2021;19(1):1–1.

