## Supplementary Material for "CPSM: R-package of an Automated Machine Learning Pipeline for Predicting the Survival Probability of Single Cancer Patient"

### 1. Description and Application of Predictive Models

#### 1.1 Model-1: Clinical Features Only

This model is constructed using significant clinical features, identified by their p-value ($p < 0.05$) through the **Univariate_sig_features_f** module of the CPSM package. The clinical features selected demonstrate strong prognostic potential for stratifying survival-risk groups. This approach is particularly effective when the available data consists solely of categorical clinical variables (e.g., age, gender, IDH1 mutation status). While this model can provide valuable insights based on clinical data alone, one challenge is that some key features may not be present for all samples, leading to incomplete coverage in certain cases.

#### 1.2 Model-2: PI Score-Based Model

This model is developed using the PI score, which is generated via the **Lasso_PI_scores_f** feature selection module of the CPSM package. It is designed to handle high-dimensional datasets, where a large number of numeric features (e.g., ~20K gene or protein expression data) are present. The PI score helps to identify and retain only the features relevant to survival outcomes from this huge pool of features, allowing the model to focus on a more refined set of predictors, thus reducing dimensionality without losing predictive power. This model is highly effective when dealing with large-scale genomic or proteomic data.

#### 1.3 Model-3: Combined Clinical and PI Score Model

This model integrates both PI scores and significant clinical features, making it suitable when both clinical and numeric (e.g., gene expression) features are available in the dataset. By combining these two types of features, the model leverages the strengths of both data types—clinical variables that are known to impact survival outcomes and high-dimensional numerical features like genes, which may provide additional prognostic information. This hybrid approach allows for more comprehensive survival analysis, especially when a large dataset with both clinical and molecular data is available.

#### 1.4 Model-4: Univariate Feature Selection for Clinical and Genomic Features

In this model, both significant clinical and gene expression features are selected through the **Univariate_sig_features_f** module of the CPSM package. This function identifies features with significant prognostic relevance in univariate survival analysis. This model is particularly useful when there is a small set of predefined genes or clinical features, for example, when a user has identified approximately 20-30 key genes through in-vitro experiments. These selected features are then assessed for their prognostic potential using the **Univariate_sig_features_f** feature selection module of CPSM, after which the model can be built using only the significant features. This approach ensures that the final model is both parsimonious and highly relevant to the survival prediction task, focusing on limited available features set in the data.

### 2. Case Studies for the CPSM package

To demonstrate the utility of the CPSM package, we conducted case studies using RNAseq gene expression datasets from TCGA for glioblastoma (GBM) and kidney Renal Clear Cell Carcinoma (KIRC). These datasets have RNA expression in terms of FPKM values along with 19 clinical and demographic features including age, gender, ajcc pathologic tumor stage etc. and 4 types of survival include OS (overall survival), Notably, in these case studies, we focused to identify prognostic features and subsequent survival probability prediction models development with respect to overall survival (OS) only.

Tables S1. Complete list of types of RNA transcripts, clinical features and survival types present in TCGA-GBM and TCGA-KIRC data used in case studies.

| RNA Transcripts in RNA | Features in Clinical data | Types of Survival in Data |
|---|---|---|
| 38,693 RNA transcripts include 19,962 protein-coding, 16,901 lncRNA, 1,881 miRNA, 5 transcribed unprocessed pseudogene, 1 transcribed processed pseudogene, 1 unprocessed pseudogene, and 1 ribozyme encoding RNA transcripts | age, subtype, gender, race, ajcc pathologic tumor stage, histological type, histological grade, treatment outcome first course, radiation treatment adjuvant, sample type (primary/recurrent). | 4 types of survival include: (1) OS (overall survival), (2) PFS (progression-free survival), (3) DSS (disease-specific survival), (4) DFS (Disease-free survival). In the data, column names OS, PFS, DSS and DFS represent event information, while OS.time, PFS.time, DSS.time and DFS.time indicate survival time in days. |

#### 2.1 Case Study1 – TCGA-GBM

For this case study, we used TCGA-GBM RNA expression data along with available clinical information. Our input data has 164 samples 38,712 features (38,693 RNA transcripts + 19 clinical and demographic features).

##### 2.1.1 Data Pre-processing and Preparation

We pre-processed our data using the *data_process_f* function of the CPSM package and got processed data with 161 samples and a new column "OS_month" to the data containing OS time in months. Then, we split data into training and test subsets using the *tr_test_f* function with a fraction of 0.9. and obtained training data with 144 samples and test data with 17 samples. Next, we performed data normalization using the *train_test_normalization_f* function of the CPSM package.

#### 2.1.2 Feature Selection

Next, we performed feature selection using the *Lasso_PI_scores_f* and the *Univariate_sig_features_f* functions of CPSM package. The *Lasso_PI_scores_f* gave us the following outputs: (1) a list of 36 features selected by LASSO with their beta coefficient values (Table S1), (2) Training and Test dataset containing the expression of selected genes and PI score (3) Lasso Regression Lambda plot (as shown in Figure S1).

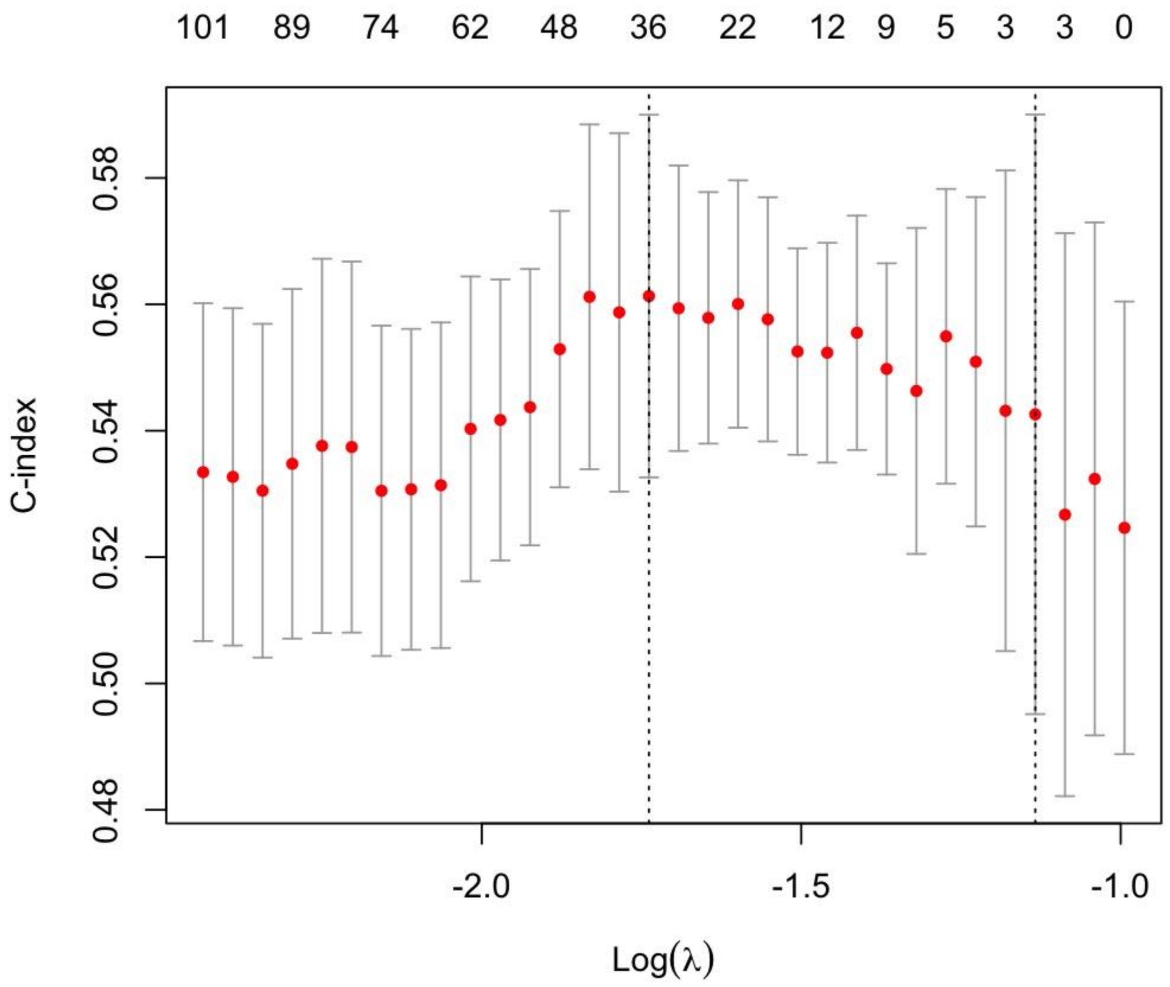


Supplementary Figure S1: Lasso Regression Lambda plot representing 36 features selected by LASSO model for TCGA-GBM data.

Next, the *Univariate_sig_features_f* gave us the following outputs: (1) a table of 2,262 features selected by univariate *Univariate_sig_features_f* function along with their corresponding coefficient values, HR value, P-values, C-Index values. (2) Training and Test datasets containing the gene expression of significant 2,262 genes.

#### 2.1.3 Model development for predicting the survival probability of patients

Following the selection of significant prognostic features through LASSO regression and univariate survival analysis, we employed the MTLR_pred_model_f function from the CPSM package to develop four distinct survival prediction models aimed at estimating patient survival probabilities. The models are as follows: (1) **Model-1**, which incorporates three significant clinical features—age, subtype, and sample type; (2) **Model-2**, which is based solely on the Prognostic Index (PI) score; (3) **Model-3**, which integrates the three clinical features with the PI score; and (4) **Model-4**, which combines 2,262 univariate gene features with the three clinical features.

Models 2 emerged as the top performer among all models evaluated on the training data, achieving a C-Index of 0.8 and IBS score of 0.068 (Table S2). Similarly, on the test dataset of 17 samples, Model-2 emerged as the top performer achieving a C-Index of 0.66 and an IBS of 0.083 (Table S2). Model 4, which utilizes a set of univariate features, also demonstrated strong performance with a C-Index of 0.63 and an IBS of 0.086. For the specific patient TCGA-41-2572-01, Model-2 predicted closest survival time to the actual time, i.e., a median survival time of 13.30 months and a mean survival time of 14.56 months and (Figure S2A). Model-4 estimated a median of 14.30 months and a mean survival time of 17.13 months and while Model-1 predicted and a median of 15.35 months and a mean of 16.65 months. Model-3, which combines clinical features and the PI score, predicted a median survival of 16.49 months and a mean survival time of 18.11 months.

Despite comparable performance between Model 2 and Model 4, the disparity in feature count is significant; Model 2 is based on only 36 RNA transcript features, whereas Model 4 utilizes 2,262 RNA transcripts.

Table S2: Performance of models on Training and Test datasets of TCGA-GBM data.

| Model | Feature Type | Dataset | IBS_Score | c_index |
|---|---|---|---|---|
| Model1 | Clinical Features | Training set | 0.091 | 0.64 |
| | | Test set | 0.092 | 0.54 |
| Model2 | PI score | Training set | 0.068 | 0.8 |
| | | Test set | 0.083 | 0.66 |
| Model3 | PI score + Clinical Features | Training set | 0.07 | 0.79 |
| | | Test set | 0.088 | 0.61 |
| Model4 | Univariate features + Clinical features | Training set | 0.077 | 0.75 |
| | | Test set | 0.092 | 0.59 |

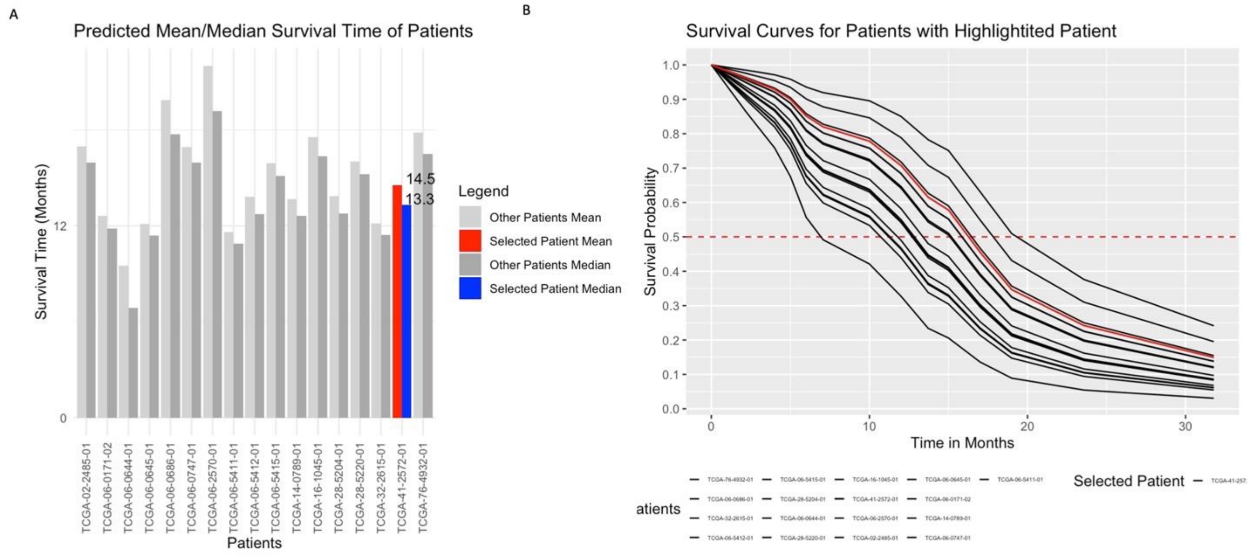


Figure S2: Prediction Results using Model2 based on PI score of TCGA-GBM test data. (A) Bar plot representing the predicted mean (red) and median (blue) survival time of selected patient (TCGA-41-2572-01) along with other samples (grey) in test data with predicted mean (light grey) and median (dark grey) survival time of patients. (B) Survival curve plot representing survival probability of selected patients, where each curve represents survival probability of individual patients over a period of time, red curve represents survival probability of selected patient (TCGA-41-2572-01) along with survival probabilities of other patients (black color). Dotted red line represent the median line with 50% survival probability.

#### 2.1.4 Biological Implication of 36 genes

Differentiating genes into protein-coding and long non-coding RNA (LincRNA) transcripts is essential for grasping their unique biological functions, regulatory mechanisms, and roles in disease, thereby providing insights into therapeutic targets and genomic organization. Among 36 features selected by the *Lasso_PI_scores_f* function, 15 are protein-coding and 21 LincRNA transcripts (as shown in Table S3). Further, 28 were identified as good prognostic factors, indicated by negative beta coefficients, while 8 were classified as bad prognostic factors, exhibiting positive beta coefficients. The gene enrichment analysis of protein coding genes using the Enrichr and STRING databases (Kuleshov, et al., 2016; Szklarczyk, et al., 2021) indicated the association of genes with Interleukin-27, Interleukin-6, Interleukin-12, Interleukin-35 Signaling pathways, RHOV GTPase Cycle pathways, transcriptional regulation, and translational initiation and elongation signaling pathway, cell development pathways, and TGF beta signaling pathways, etc.

The survival unfavorable genes can be further explored for their drug-target potential to improve the prognosis of patient.

Table S3. A list of 36 features (along with beta coefficient values and transcript type) selected by the LASSO Regression for TCGA-GBM data based on which PI score generated.

| **Gene** | **Beta Coeff** | **Gene Type** |
|---|---|---|
| *IL27RA* | 0.047 | Protein-coding gene |
| *BMPR1A* | -0.158 | Protein-coding gene |
| *NSUN5* | 0.009 | Protein-coding gene |
| *MED10* | 0.092 | Protein-coding gene |
| *ZNF189* | -0.141 | Protein-coding gene |
| *OSMR* | 0.002 | Protein-coding gene |
| *ZIC3* | -0.364 | Protein-coding gene |
| *RPL39L* | 0.05 | Protein-coding gene |
| *FBXW12* | -0.092 | Protein-coding gene |
| *ARHGAP12* | -0.042 | Protein-coding gene |
| *HOXB2* | 0.04 | Protein-coding gene |
| *ZNF732* | -0.254 | Protein-coding gene |
| *SPOUT1* | -0.112 | Protein-coding gene |
| *RUFY2* | -0.135 | Protein-coding gene |
| *LGALS8-AS1* | 1.019 | Long-non coding RNA |
| *SOX21-AS1* | -0.135 | Long-non coding RNA |
| *SMAD9- IT1* | -0.095 | Long-non coding RNA |
| *MLIP.IT1* | -0.205 | Long-non coding RNA |
| *AC092894.1* | -3.412 | Long-non coding RNA |
| *LINC02108* | -5.642 | Long-non coding RNA |
| *AC093766.1* | -0.871 | Long-non coding RNA |
| *AC136475.4* | -0.728 | Long-non coding RNA |
| *AC021351.1* | -0.164 | Long-non coding RNA |
| *AC013452.2* | -0.078 | Long-non coding RNA |
| *RARA.AS1* | 0.011 | Long-non coding RNA |

| | | |
|---|---|---|
| *AC091132.4* | -0.15 | Long-non coding RNA |
| *AC009686.2* | -0.027 | Long-non coding RNA |
| *AL121992.3* | -0.442 | Long-non coding RNA |
| *AC067750.1* | -0.007 | Long-non coding RNA |
| *MIR6862.1* | -0.505 | Long-non coding RNA |
| *AC099671.1* | -0.44 | Long-non coding RNA |
| *AL441992.2* | -0.072 | Protein-coding gene |
| *AC073968.2* | -0.228 | Long-non coding RNA |
| *AL391538.1* | -0.639 | Long-non coding RNA |
| *AC005702.5* | -0.331 | Long-non coding RNA |
| *AC100801.2* | -0.378 | Long-non coding RNA |

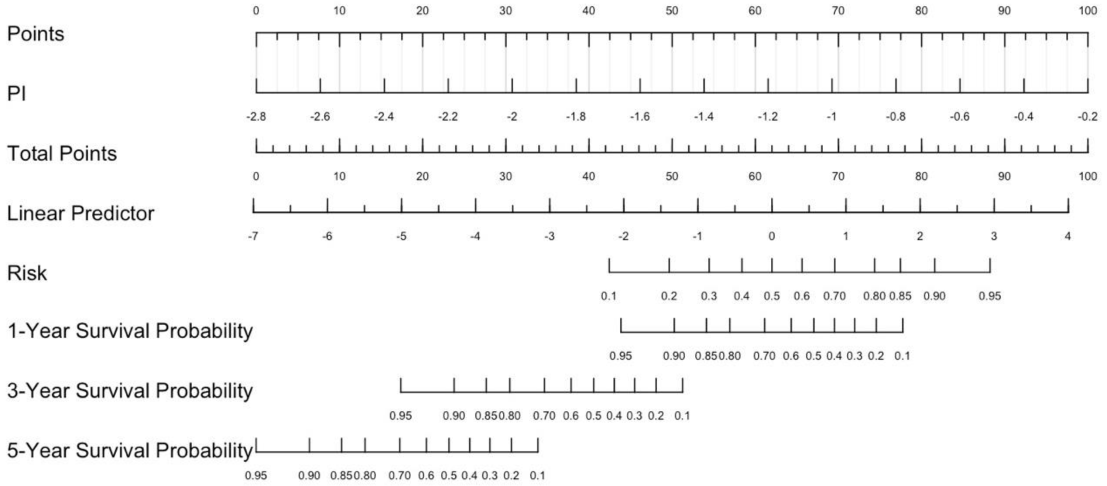


Figure S3: Nomogram for TCGA-GBM data using based on PI score to predict risk, 1,3,5-year survival of patients.

### 2.2 Case Study 2 – TCGA-KIRC

In this case study, we aimed to identify features that predict the survival probability of patients with Kidney Renal Clear Cell Carcinoma (KIRC) using the CPSM package and RNA expression data from TCGA-KIRC. The dataset comprises 534 samples and 38,712 features

#### 2.2.1 Data Pre-processing and Preparation

Following the methodology in Case Study 1, we employed the **data_process_f** function of the CPSM package to process the KIRC data, resulting in a dataset with 526 samples after removing 8 samples due to missing overall survival (OS) information. Additionally, a new column titled "OS_month" was introduced, representing OS time in months. Subsequently, we partitioned the data into training and test sets in a 90:10 ratio using the **tr_test_f** function, yielding 473 samples for training and 53 samples for testing. As in Case Study 1, we applied quantile normalization to the gene expression data via the **train_test_normalization_f** function of the CPSM package.

#### 2.2.2 Feature Selection

Next, we selected survival-relevant features for the TCGA-KIRC dataset using the **Lasso_PI_scores_f and Univariate_sig_features_f** functions from the CPSM package. The Lasso_PI_scores_f function yielded the following results: (1) a list of 9 genes identified by LASSO, along with their corresponding beta coefficient values; (2) training and test datasets containing the gene expression levels of these 9 genes, along with the predicted index (PI) score; and (3) a Lasso Regression lambda plot (Figure S4).

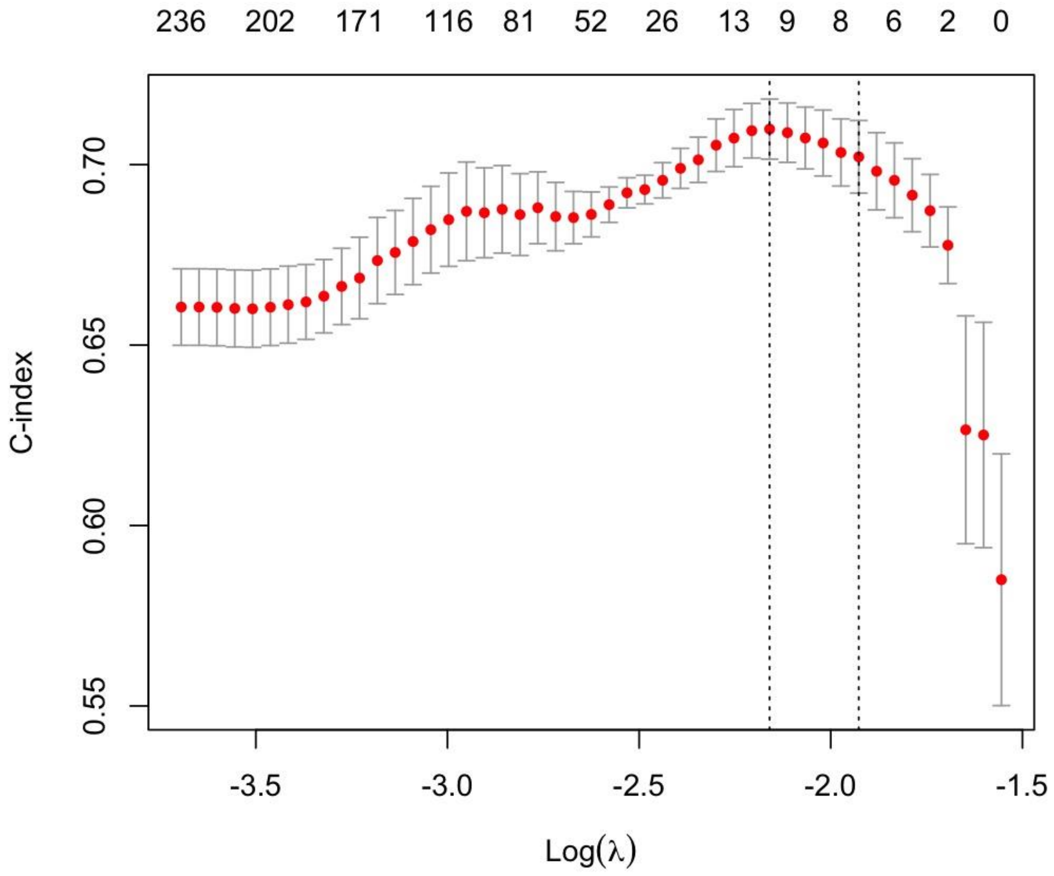


Supplementary Figure S4: Lasso Regression Lambda plot for TCGA-KIRC data representing 9 features selected by LASSO model.

The Univariate_sig_features_f function provided the following outputs: (1) a table listing 11,319 features along with their corresponding hazard ratio (HR) values, p-values, and concordance index (C-Index) values; and (2) training and test datasets containing the gene expression levels of these 11,319 significant genes.

#### 2.2.3 Model development for predicting the survival probability of patients

Next, we developed four survival probability prediction model to predict the survival probability of patients include (1) Model-1 based on 3 significant clinical features (age, subtype, histological grade), (2) Model-2 using the PI score, (3) Model3 combining 3 clinical features with PI score, (4) combining 11,319 univariate genes with 3 clinical features. Among the evaluated models for this study, Model 3 demonstrated the highest performance on the training dataset, achieving a C-Index of 0.80 and an Integrated Brier Score (IBS) of 0.79 (Table S4). In the subsequent analysis of the full test dataset of 53 samples, Model 3, continued to lead with a C-Index of 0.76 and an IBS of 0.23. Model-1, which utilizes three clinical features, also exhibited commendable performance with a C-Index of 0.73 and an IBS of 0.25 (Table S4).

In the case of specific patient **TCGA-CJ-6030-01**, who had an overall survival (OS) time of 76 months, Model 3 estimated a median survival time of 73.81 months and a mean survival time of 84.85 months (Figure S5A). In comparison, Model 1 predicted a median survival time of 80.10 months and a mean of 90.40 months; while Model 2 provided estimates of 83.60 months for median and 92.19 months for mean survival time. Notably, Model 4 demonstrated significantly inferior performance, predicting a median survival time of 31.09 months and a mean survival time of 60.32 months.

Table S4: Performance of models on Training and Test datasets of TCGA-KIRC data.

| Model | Feature Type | Dataset | IBS_Score | c_index |
|---|---|---|---|---|
| Model1 | Clinical Features | Training set | 0.197 | 0.71 |
| | | Test set | 0.25 | 0.73 |
| Model2 | PI score | Training set | 0.177 | 0.75 |
| | | Test set | 0.297 | 0.70 |
| Model3 | PI score + Clinical Features | Training set | 0.16 | 0.79 |
| | | Test set | 0.23 | 0.76 |
| Model5 | Univariate features + Clinical features | Training set | 0.24 | 0.70 |
| | | Test set | 0.295 | 0.67 |

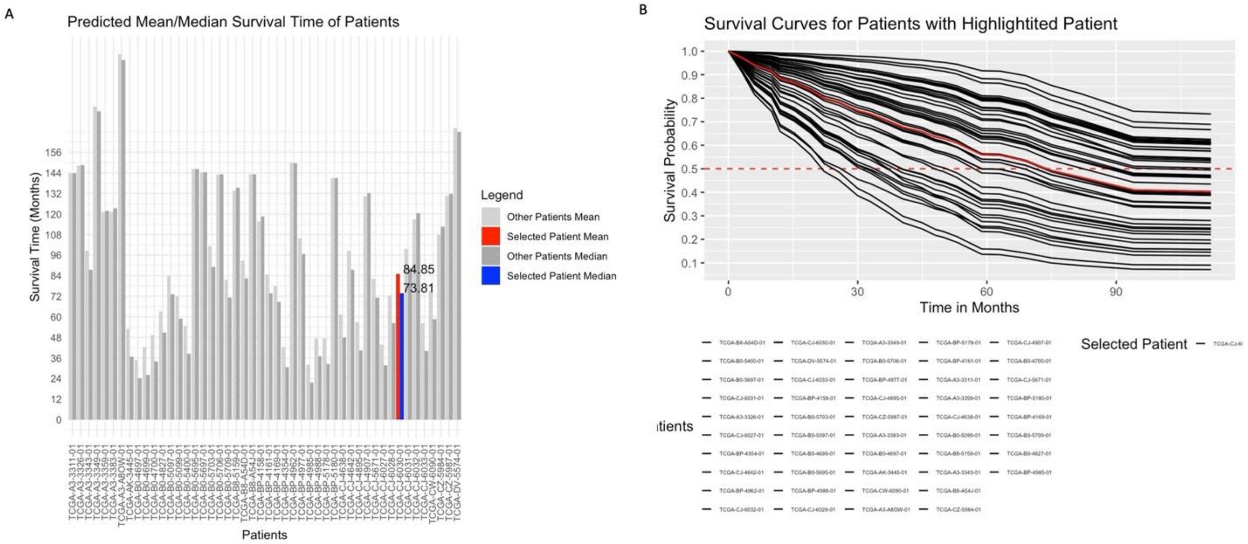

Figure S5: Prediction Results using model3 based on PI score with clinical features of TCGA-KIRC test data. (A) Bar plot representing the predicted mean (red) and median (blue) survival time of selected patient (**TCGA-CJ-6030-01**) along with other samples (grey) in test data with predicted mean (light grey) and median (dark grey) survival time of patients. (B) Survival curve plot representing survival probability of selected patients, where each curve represents survival probability of individual patients over a period of time, red curve represents survival probability of selected patient (**TCGA-CJ-6030-01)** along with survival probabilities of other patients (black color). Dotted red line represent the median line with 50% survival probability.

#### 2.2.4 Biological Implication of 9 RNA features

Among the nine selected features, five are protein-coding genes: ***NARF***, ***SLC16A12***, ***SORBS2***, ***DONSON***, and ***SOWAHB***. The remaining four are long intergenic non-coding RNAs (lincRNAs): **AL357140.2**, **LINC01605**, **DLGAP1-AS2**, and **AL355999.1** (Table S5). Further, 4 genes can be categorized as good prognostic factors (negative beta coefficients), while 5 were deemed bad prognostic factors (positive beta coefficients) (Table S5). Gene enrichment analysis conducted using the Enrichr database and STRING (Kuleshov, et al., 2016; Szklarczyk, et al., 2021) revealed associations of these protein-coding genes with several biological processes, including the Mitotic G2/M Transition Checkpoint, DNA Integrity Checkpoint Signaling, Mitotic DNA Damage Checkpoint Signaling, Carboxylic Acid Transport, and Vascular Transport. The identification of unfavorable survival genes presents an opportunity for further exploration in in vitro and in vivo studies to assess their potential as drug targets aimed at improving patient prognosis.

Table S5. A list of 9 features (along with beta coefficient values and transcript type) selected by the LASSO Regression for TCGA-KIRC data based on which PI score generated.

| **Gene** | **Beta Coeff** | **Transcript Type** |
|---|---|---|
| *NARF* | 0.099 | Protein coding |
| *SLC16A12* | -0.031 | Protein coding |
| *SORBS2* | -0.172 | Protein coding |
| *DONSON* | 0.103 | Protein coding |
| *SOWAHB* | -0.122 | Protein coding |
| *AL357140.2* | -0.35 | lncRNA |
| *LINC01605* | 0.084 | lncRNA |
| *DLGAP1-AS2* | 0.106 | lncRNA |
| *AL355999.1* | 0.148 | lncRNA |

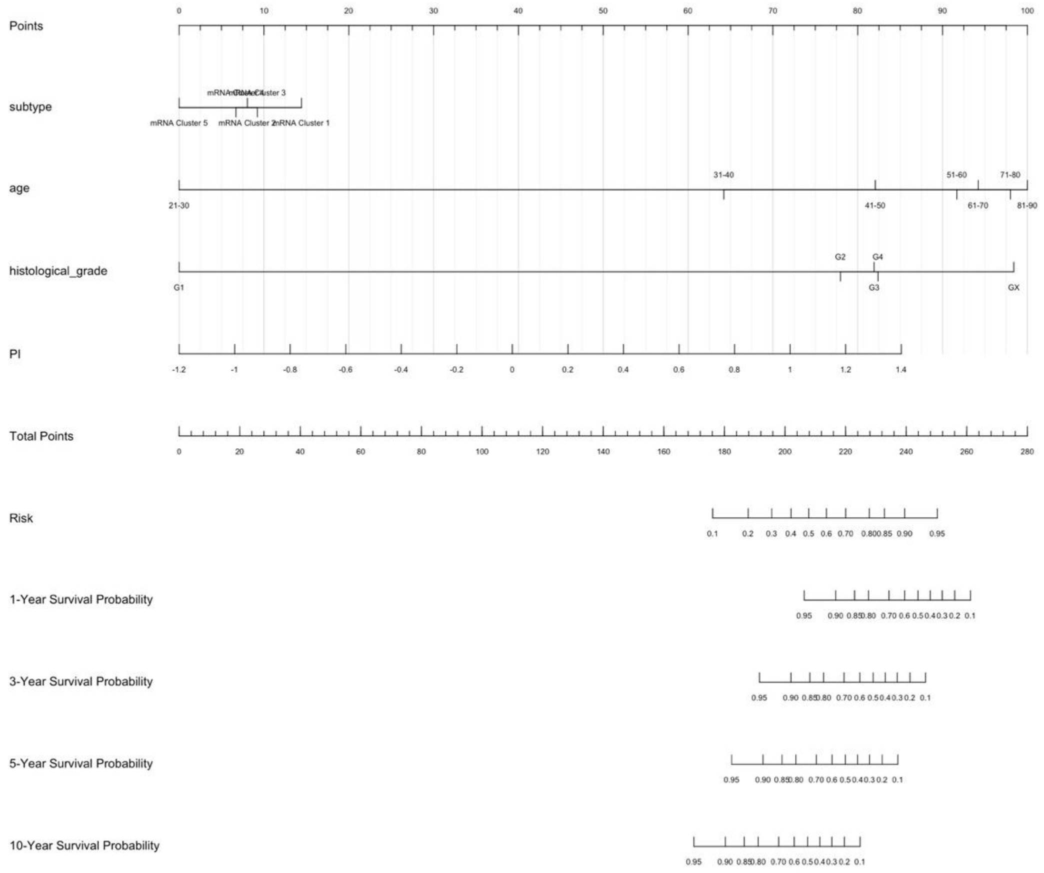


Figure S6: Nomogram for TCGA-KIRC data using based on clinical features and PI score to predict risk, 1,3,5, and 10-year survival of patients.

### References

Kuleshov, M.V., *et al*. Enrichr: a comprehensive gene set enrichment analysis web server 2016 update. *Nucleic Acids Res* 2016;44(W1):W90-97.

Szklarczyk, D., *et al*. The STRING database in 2021: customizable protein-protein networks, and functional characterization of user-uploaded gene/measurement sets. *Nucleic Acids Res* 2021;49(D1):D605-D612.
